# Immunopharmacological evaluation of adjuvant efficacy of Monophosphoryl lipid-A and CpG ODN with SARS-CoV-2 RBD antigen

**DOI:** 10.1101/2022.04.04.486920

**Authors:** Shainy Sambyal, Vemireddy Sravanthi, Ritika Khatri, Gazala Siddqui, Shubbir Ahmed, Sweety Samal, Halmuthur M Sampath Kumar

**Author notes:** **Corresponding author:** Halmuthur M. Sampath Kumar, Vaccine Immunology Laboratory, OSPC Division, CSIR-Indian Institute of Chemical Technology, Hyderabad - 500007, India. **Email:**.

## Abstract

SARS-CoV-2 infection has made the mankind to witness most sever and serious pandemic situation in the history. Millions of people have suffered and are still suffering with this infection which has caused a mass mortality in the past three years. Development of an effective vaccine to control the spread of infection and to prevent this viral infection is need of the hour. Adjuvanted vaccines have proven their efficacy in controlling many other viral infections like flu, keeping this context in view we have evaluated the immunopharmacological efficacy of two adjuvants MPL-A and CpG ODN in combination with MF59 emulsion against SARS-CoV-2 antigen. From the data obtained we can infer that both the adjuvants were capable of eliciting a potent antibody response against antigen alone and MF59 groups. Comparatively MPL-A was eliciting a Th1 polarized response in terms of IgG2a and cytokine production. Both the adjuvants were capable of enhancing the CD 4, 8 and 19 cell populations. Overall the pre clinical evaluation has given a clue of the effectiveness of MPL-A and CpG adjuvants against SARS-CoV-2 antigen.

## Introduction

Corona virus belongs to the family of *Coronaviridae* which majorly infect mammals, amphibians and birds. This group of viruses can cause mild to severe respiratory tract infections in humans. Between 2002-2012 highly infectious viruses (SARS-CoV and MERS-CoV) with zoonotic origin caused lethal respiratory illness in humans leading to new public health apprehension in the twenty first century [1]. Recently, since past two-three years novel coronavirus namely SARS-CoV-2, reportedly emerged from Wuhan city, China caused a pandemic situation throughout the world and has mutated to other virulent forms leading to mass mortality within no time [2].

As of time being, there are no certified treatments; prevention plays a very crucial role in controlling the infection [3]. Many factors like, infectivity yet before the onset of warning sign in incubation period, variation in incubation period, rapid transmission from asymptomatic people, extended duration of the transmission even after clinical recovery *etc* make prevention very difficult [4]. Research is centered towards developing an effective and efficient vaccine that can combat this infection in particular with emerging mutation strains. Role of adjuvants in developing such vaccines cannot be denied and many of the present day vaccines like Covxin (TLR 7/8 ligand) have an adjuvant incorporated within them [5]. The research carried out by various people from across the globe stress on the need of using an adjuvant for successful development of a vaccine which would be capable of eliciting a protective response against wide varieties of mutants that are evolving.

Adjuvant use in vaccine is an age old phenomena and effectiveness of adjuvanted vaccine for various diseases like flu has been well established. Adjuvants such as MF59, AS03, MPL-A, CpG, and QS-21 etc have been widely applied for human used and were proven safe [6]. Use of these human approved adjuvants for SARS-CoV-2 vaccine development can be effective and time saving in this pandemic situation.

In this study, we want to investigate that immunological effect of adjuvants that would generate a strong humoral response against SARS-CoV-2 formulated with O/W based delivery system - MF59. In order to study this we have formulated SARS-CoV-2 Spike Glycoprotein (S1) and MF59 alone or in combination with Th1 adjuvants at varying concentration of 10, 20 and 30μg. Both the standard adjuvants (MPL-A and CpG ODN) tested are commercially available and used in several FDA licensed vaccines. Monophosphoryl lipid A (MPL-A) a TLR-4 vaccine adjuvant synthesized by hydrolysis of native diphosphoryl lipid A, the immunoreactive component of Lipopolysaccharide (LPS) present on the lipid bilayer of *Salmonella Minnesota* [24]. Cytosine-guanine oligodeoxynucleotide (CpG ODN) is a TLR-9 agonist bacterial DNA motifs and an effective catalyst of innate immunity.

## 2. Materials and methods

### 2.1 Chemicals and reagents

SARS-CoV-2 Spike Glycoprotein (S1) RBD- (amino acids 328 to 531) (THSTI, India), MPL-A (InvivoGen, USA), CpG ODN (InvivoGen, USA), RPMI 1640 (HiMedia, Mumbai), DPBS (HiMedia, Mumbai), sodium bicarbonate (Sigma-Aldrich, USA), sodium carbonate (Sigma-Aldrich, USA), Trypan blue (HiMedia, Mumbai), Thiazolyl blue tetrazolium bromide i.e. MTT (Sigma-Aldrich, USA), anti mouse cytokines for ELISA-IFN-gamma, TNF-alpha, IL-6, IL-1β (BioLegend, Switzerland), flurochrome conjugated FACS anti mouse antibodies-CD16/32, CD4, CD8, CD19 (BD, USA), TMB substrate (BD, USA), HRP streptavidin (BioLegend, Switzerland), HRP Goat anti-mouse IgG antibody (BioLegend, Switzerland), anti-mouse IgG1 horseradish peroxidase conjugates (BioLegend, Switzerland), anti-mouse IgG2a horseradish peroxidase conjugates (BioLegend, Switzerland) were used.

### 2.2 Methodology

#### 2.2.1 Immunization

4-6 weeks female BALB/c mice purchased from RCC Laboratories India Private Limited were grouped as antigen alone (1μg/dose), MF59 either in combination with antigen alone or in combination with MPL-A and CpG ODN at varying concentration of 10, 20 and 30μg. All the mice were injected subcutaneously on 0^th^ day and a booster was given on 14^th^ day under the guidelines of IAEC with IAEC No. *IICT-IAEC-025-2020*. Retro-orbital blood and spleen was collected on 28^th^ day.

#### 2.2.2 Sera collection

The retro-orbital blood collected from each mice on 28th day was centrifuged at 2000rpm for 20 min and supernatant was collected to fresh tubes separately. The sera was stored at 0 C and later was used for antibody titer: IgG, IgG1, IgG2a.

#### 2.2.3 Spleen and splenocyte isolation

Mice were sacrificed with light ether anesthesia. The spleen were collected in ice chilled PBS from each group separately. In laboratory conditions the spleen were homozinised in RPMI complete media using 70μm cell strainer. The cells were treated with ACK lysis buffer (0.5M ammonium chloride, 10mM potassium bicarbonate and 0.1mM disodium ethylene diamine tetraacetic acid, pH 7.2) to lyse RBC’s i.e. 1ml/spleen for 1min 30sec. Lymphocytes so collected were then washed with PBS and cell frequency was calculated by opting trypan blue method (Hemocytometer).

#### 2.2.4 Estimation of IgG antibody

The sera collected from retro-orbital blood of immunized mice was analysed for quantigication of IgG response via indirect ELISA. Briefly, SARS-CoV-2 antigen diluted in bicarbonate-carbonate buffer were coated on 96 well plate at 4°C for overnight. Post incubation the plates were washed with PBST (PBS containing 0.05% (v/v) Tween 20) and blocked with 1% BSA for 1 hour at RT. After three wash, serially diluted sera sample was added and kept for incubation at 37°C for next two hours with 5% CO_2_. 100μl of anti-mouse IgG horseradish peroxidase conjugates diluted 1:3000 was added and kept undisturbed for 1 hour at RT in dark. The plates after incubation were added with 100μl of TMB substrate solution (1:1, Reagent A and Reagent B) and after 15 min incubation development of blue color was observed. 50 μl/well of 2N H_2_SO_4_ was added to terminate the enzyme reaction. The optical density of color developed was measured in Tecan Multimode reader at 450nm [7].

#### 2.2.5 Estimation of IgG1 and IgG2a subtypes

Similar to that of IgG antibody titer, IgG1 and IgG2a subtypes titer was performed. The plates were coated with antigen, blocked with 1% BSA, addition of serially diluted serum sample tagged with 1:3000 diluted anti-mouse IgG1 horseradish peroxidase conjugates and anti-mouse IgG2a horseradish peroxidase conjugates. Color was developed with TMB and absorbance was determined opting same Tecan multimode reader at 450nm [7].

#### 2.2.6 Estimation of CD4, CD8 and CD19 markers

CD4, CD8 and CD19 population was evaluated using FACS Verse technology. Briefly, 3X10^5^splenocytes were elicited in eppendorf tubes separately as per groups. The cells were washed with 500μl of FACS staining buffer. To avoid non-specific binding of Fc receptors chiefly present on B-cells, NK cells, macrophages, dendritic cells with antibodies via their consistent domain, the Fc receptors were blocked with CD16/32 for 15 min at 4°C. Again cells were washed with 250μl of FACS staining buffer to clear away unbound antibodies. The cells were then stained with 5μl of PE-antiCD4, FITC-A antiCD8 and FITC-A antiCD19 flurochrome tagged antibodies for 45 min at 4°C. The pellets so collected were resuspended in sheath fluid and cell population was quantified using BD FACSsuit software [8].

#### 2.2.7 Splenocyte proliferation

Splenocytes from selected group were seeded in 96 well plates with cell count 1X10^5^cells/well. The cells were restimulated by using SARS-COV-2 antigen, LPS and CON A with concentration 0.1, 10 and 2μg/ml respectively. The plates were incubated for 48 hours at 37°C having 5% CO_2_. After the fulfillment of incubation, 20μl/well MTT reagent (5mg/ml PBS) was added and incubated at 37°C for next 3 hours. Post incubation the untransformed MTT was replaced by 50μl of DMSO and kept for incubation at RT for about 15 min. % proliferation of the cells was calculated by measuring absorbance at 630nm and keeping cell control as reference [9].

#### 2.2.8 Estimation of Th1 and Th2 cytokines

Splenocytes from selected group were seeded in 6 well plates with cell count 1X10^5^cells/well and restimulated with 0.1μg/ml SARS-CoV-2 antigen for 48 hours at 37 °C with 5% CO_2_. The cell supernatant was quantified for cytokine estimation by imposing sandwich ELISA. Briefly, IFN-gamma, TNF-alpha and IL-6 coating antibody was diluted in bicarbonate-carbonate buffer and coated on ELISA plate for overnight at 4°C. Post incubation the plates were washed with PBST (PBS containing 0.05% (v/v) Tween 20) and blocked with 1% BSA for 1 hour at RT. After three wash, serially diluted recombinant antibody along with supernatant sample was added and ept for incubation at 37°C for next three hours with 5% CO_2_. Four times washing with PBST 20 and 100μl of detection antibody diluted in 1%BSA was added and kept at RT for 1 hour. 100μl of anti-mouse horseradish peroxidase conjugates diluted 1:3000 was added and kept undisturbed for 30min at RT in dark. The plates after incubation were added with 100μl of TMB substrate solution (1:1, Reagent A and Reagent B) and after 15 min incubation development of blue color was observed. 50 μL/well of 2N H_2_SO_4_ was added to terminate the enzyme reaction. The optical density of color developed was measured in Tecan Multimode reader at 450nm [10].

## 3. Results

### 3.1 Antibody titer

Immunized mice blood serum was used for antibody (IgG) quantification. All the immunized groups with varying concentration of 10, 20 and 30μg were analyzed and among all CpG ODN at 30μg and MPL-A at 20μg showed increased response when compared to antigen alone *i.e*. 15.23 and 13.95 folds increase respectively. In contrary when compared to MF59 it was around 2 folds. From Figure-1 we can infer that both the adjuvants at any immunized concentration the IgG response was spiking up giving a clear indication that antigen in combination with MF59 and adjuvant gives more potential response when compared to antigen alone with MF59.

**Figure-1:**
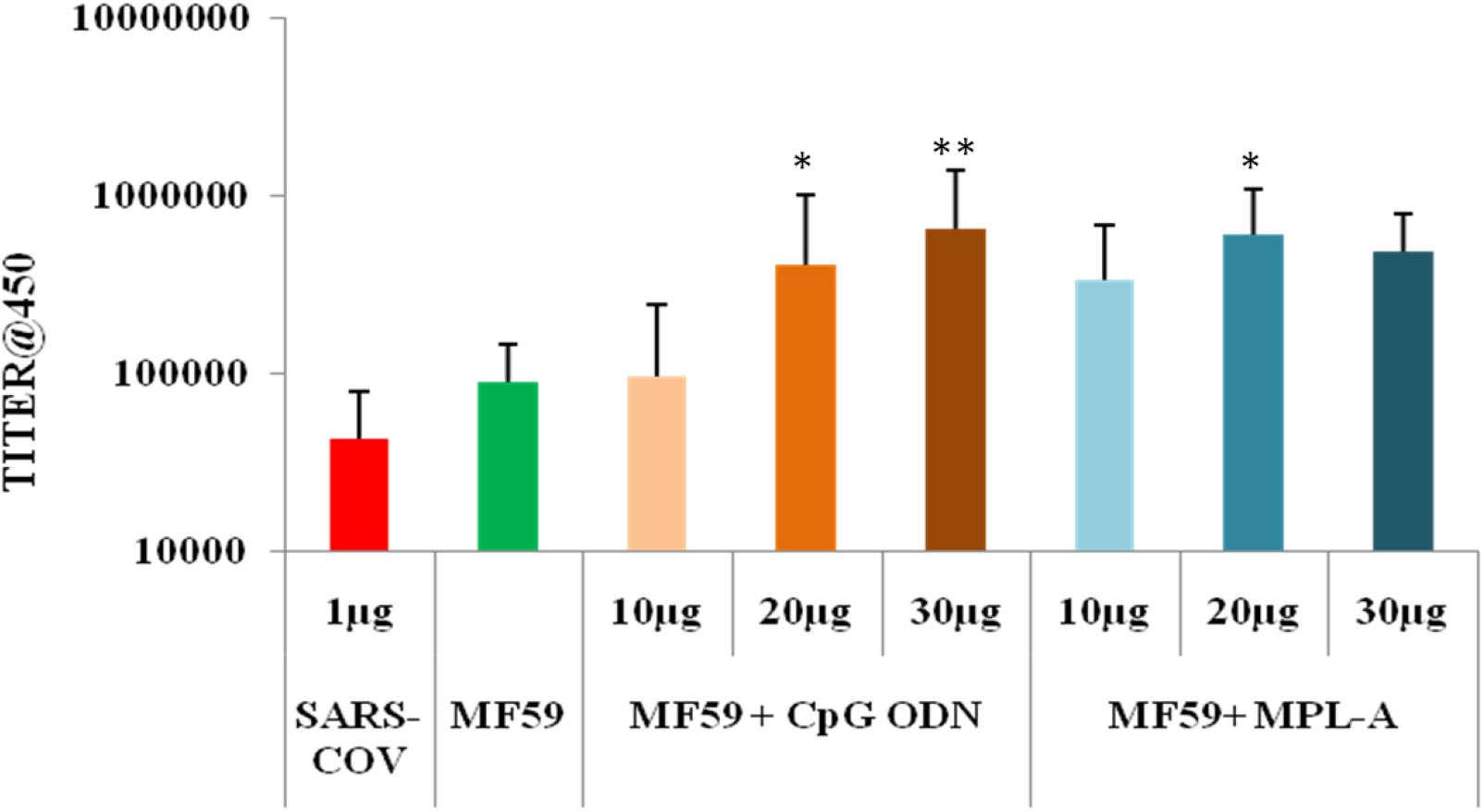
IgG titers for imuunized groups-Indirect ELISA was used to quantify the IgG antibody produced upon immunization of emulsion with adjuvants like CpG and MPL-A at various concentrations. The graphs here represent titers of mean±SD obtained from five individual mice. Student T test was profomed to determine the statistical significance of titers compared to antigen alone group wherein, *= p≤0.05 and **=p≤0.005.

### 3.2 IgG Subtypes

Similarly the groups with significant IgG titer were analyzed for subtypes: IgG1 and IgG2a. From results shown in figure 2, all the adjuvant treated groups showed higher response as compared to controls. Also IgG2a titer has spiked more than IgG1 which indicates that these adjuvanted groups were heading towards Th1 immunity where IgG2a/IgG1 >1.

**Figure-2:**
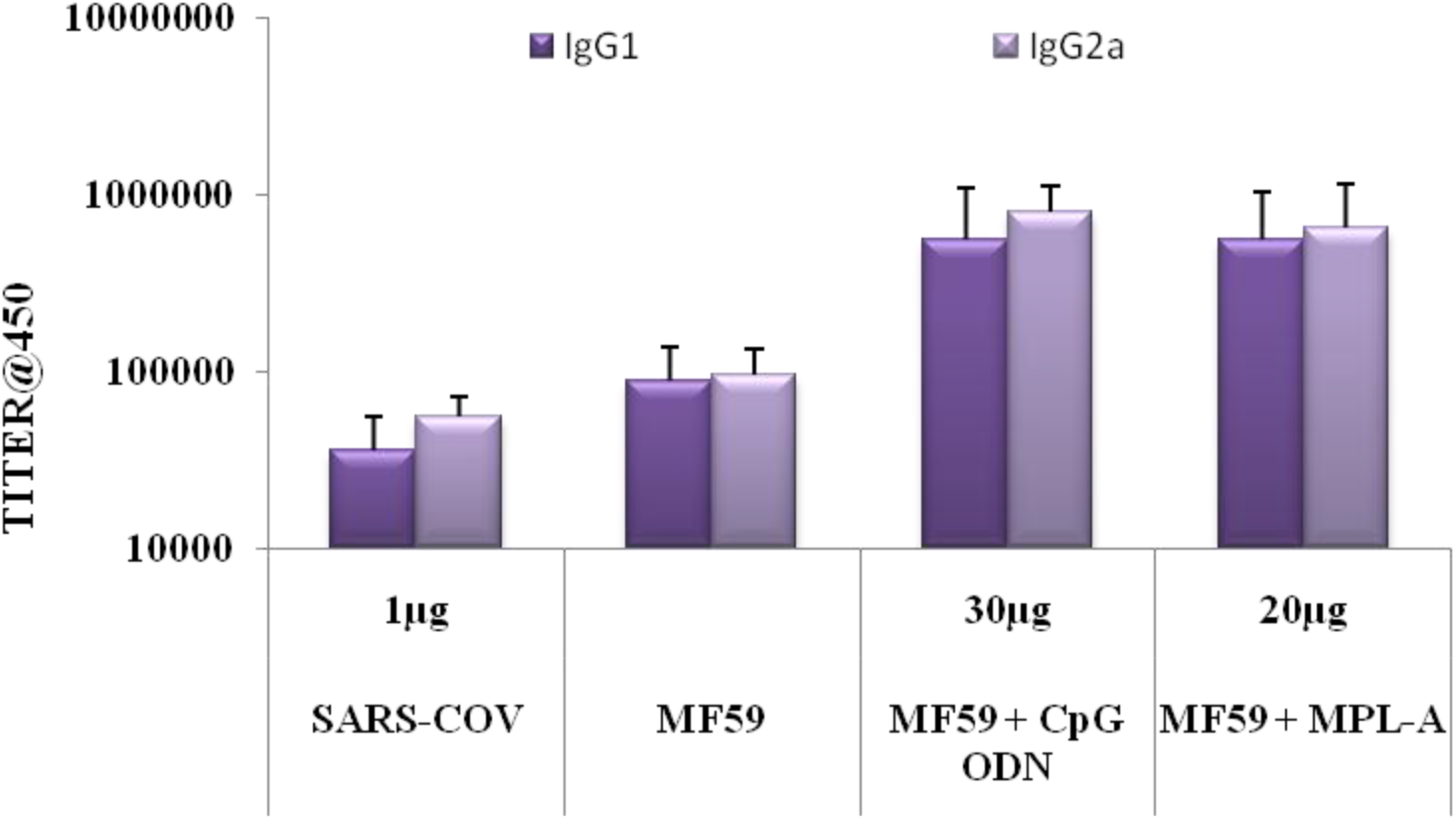
Isotype titers of adjuvanted groups-Indirect ELISA was used to quantify the IgG1 and 2a antibody produced upon immunization of emulsion with adjuvants like CpG and MPL-A at various concentrations.

### 3.3 Immunophenotyping

CD4, CD8 and CD19 surface markers were quantified for determining T cell and B cell population. From figure -3 we can infer that MPL-A 20 μg showed increased in CD4 and CD8 percent population compared to both antigen and MF59 group. Whereas MPL-A has elicited an enhancement in CD19 population too. But overall %population showed by all the groups were more than antigen alone group and more or less equal to MF59 group.

**Figure 3:**
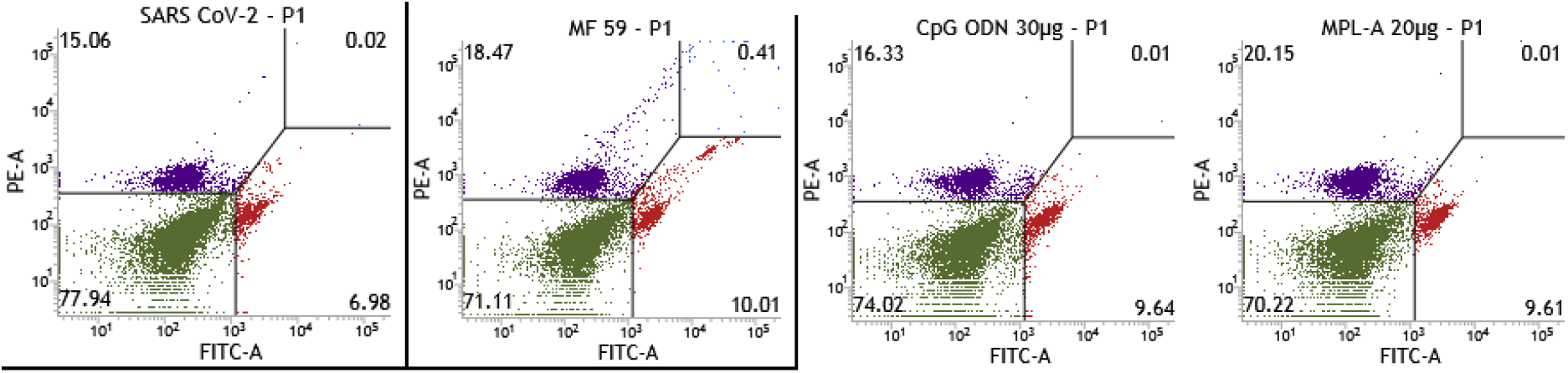
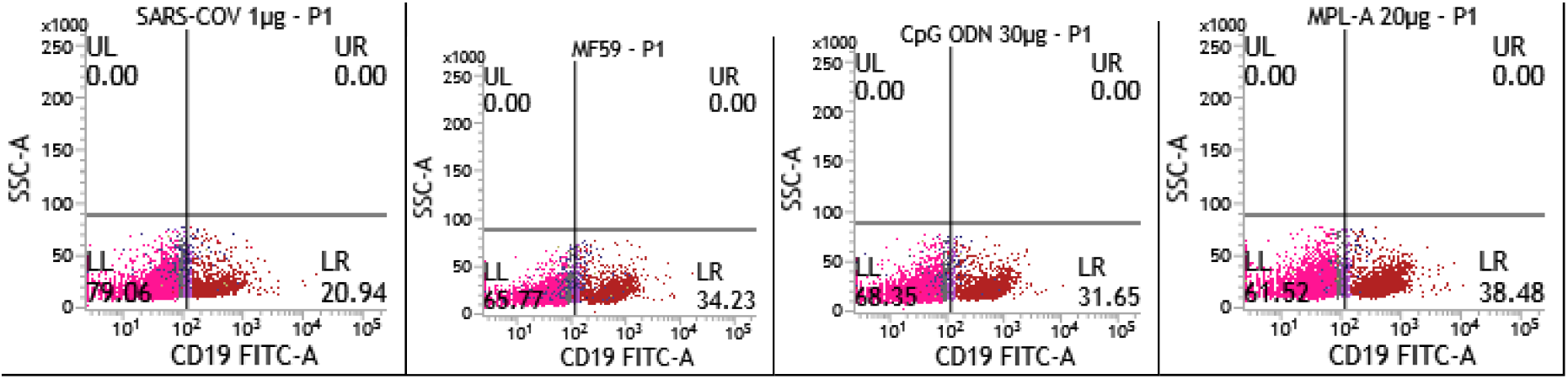
CD4, CD 8 and CD19 expression on lymphocytes-Flowcytometry was used to determine the percentage population of CD4, 8 and 19 cells from the immunized mice splenocytes.

### 3.4 Splenocyte proliferation

After 48 hours of restimulation with antigen, LPS and CON A, the %proliferation was calculated by keeping cell control as reference. From figure-4 we can conclude that among all the groups CpG ODN and MPL-A showed proliferation in LPS and CON A population where LPS represents B lymphocytes and CON A represents T lymphocytes. But overall none of the population in going below cell control which proves that none of them was cytotoxic to splenocytes.

**Figure 4:**
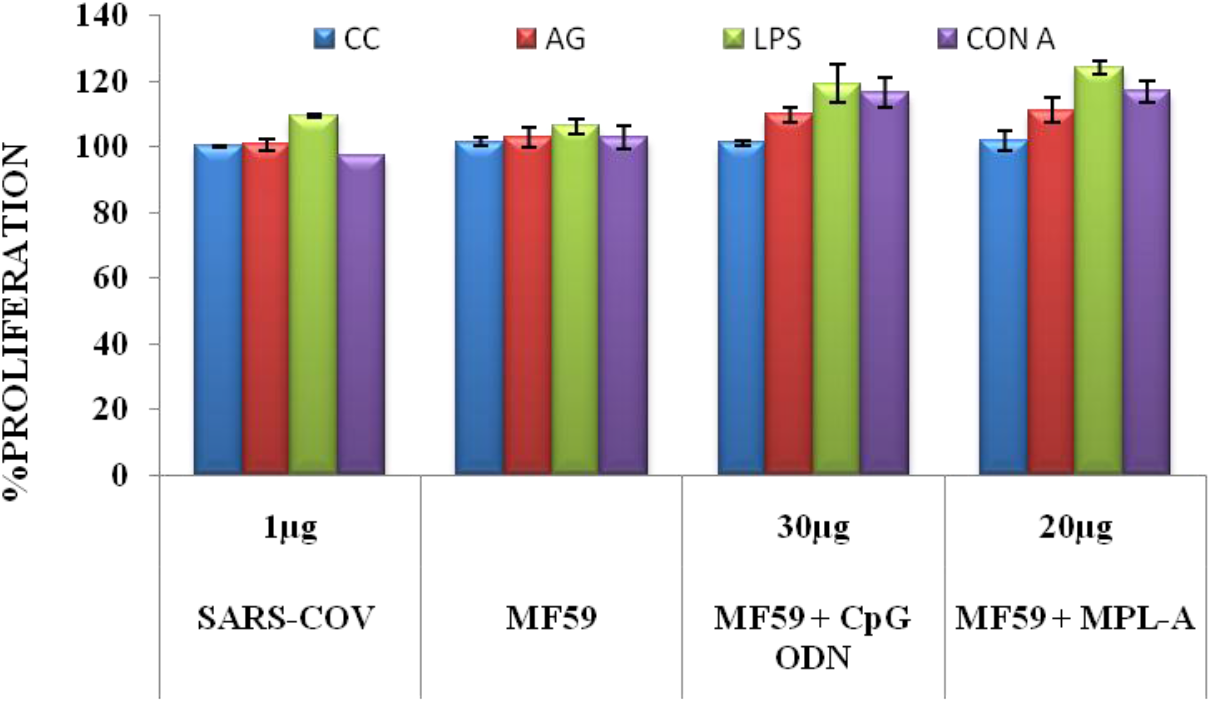
Splenocyte proliferation of immunized groups restimulated with LPS and Con A

### 3.5 Cytokine profiling

Both Th1 (IFN-gamma and TNF-alpha) and Th2 (IL-6) cytokines were profiled from the select groups along with pyrogenic cytokine IL-1β. From figure 5 we can understand that MPL-A had significant Th1 cytokine production in comparision to antigen alone group.

**Figure 5:**
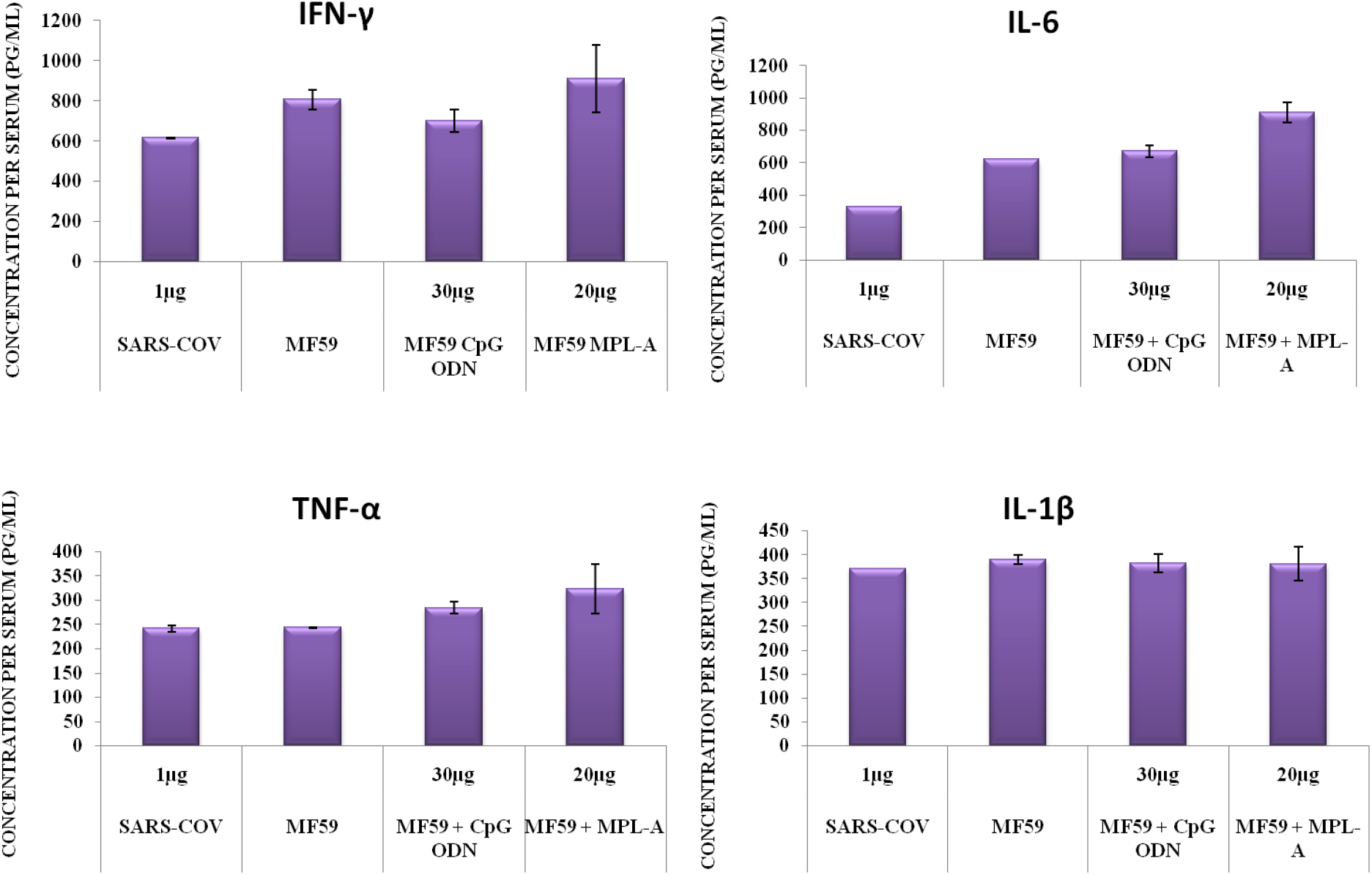
Cytokine profiles of immunized groups-Cell supernatant was used to quantify cytokines and graphs represent means±SD from three different experiments.

## Discussion

The overall aim behind this study was to find out a suitable adjuvanted based effective vaccine against SARS-CoV-2 antigen which had severe virulence world wide in no time and caused mass hospitalization and mortality to human life. The idea behind this reserach was to generate comparative study data with two standard adjuvants - CpG ODN and MPL-A in preclinical aspects tested on BALB/c mice which may be a powerful information for the world to work further on clinical trails to develop a powerful vaccine to fight COVID-19. The study behind with antibody titre *i.e*. IgG titre because immunoglobulin G cover 75% of serum antibodies in human and most common types of antibody found in human blood [11]. To fight against any viral infection the body should produce enough amount of IgG antibody. All the adjuvants tested in combination with MF59 and SARS-CoV-2 spikes the response to significant extent. Not only IgG they also proved their potential by spiking up IgG1 and IgG2a antibodies. IgG1 represents Th2 immunity and IgG2 represents Th1 immunity. IgG1 is the most abundant subclass of IgG in human blood sera and importantly mediates the antibody responses against viral pathogens. IgG2a is a predominant isotope of IgG produced in response to infection with DNA or RNA viruses. IgG2a/IgG1 was greater than 1 for all groups which heads toward Th1 immunity. Th1 immunity was preferentially regulated by IFN-gamma which helps against intracellular pathogens, such as viral infections and tumors.

The T cell immunity was regulated and monitored by CD4 and CD8 receptors present on Th cells and Tc cells respectively. Similarly activated B cells were reported by CD19 receptors. To generate a potent humoral and cell mediated immune response there should be balance between Th cells and Tc cells. Cluster of differentiation 4 (CD4) is a glycoprotein found on the surface markers present on Th cells also on macrophages and monocytes. CD4 is a co-receptor of the T cell receptor and assists the latter in communicating with antigen-presenting cells by binding to distinct regions of the antigen presenting MHC-II molecule. They do not neutralise the infections but rather trigger the body’s response to infections. CD8 is a transmembrane glycoprotein serves as co-receptor to the t cell receptors and recognise the peptides presented by MHC-I molecules.

CD8 plays role in T cell signaling and aiding with cytotoxic T cell antigen interactions. CD19 is a cell surface glycoprotein expressed by B lymphocytes and is critically involved in establishing intrinsic B cell signaling. Splenocyte proliferation was another aspect to determine the potency of adjuvants as an effective molecule against SARS-CoV-2 antigen. All the cells restimulated with LPS and CON A to determine the B lymphocytes and T lymphocytes. B cells are primarily responsible for humoral immunity (antibody mediated immunity) and T cells are involved in cell-mediated immunity. These humoral and cell mediated immunities were regulated by cytokines such as IFN-gamma, TNF-alpha and IL-6. IFN-gamma cytokine is critical to both innate and adaptive immunity and functions as a primary activator of macrophages, stimulating natural killer cells and neutrophils.

## Conclusion

The main objective of the present work is to determine the suitable adjuvant in human use for SARS-CoV-2 vaccine development in combination with MF59. From the above data we can conclude that both the adjuvants MPL-A and CpG at various concentrations had a profund impact on enhancement of antibody titer-humoral immunity along with CD4 and 8 T cell response-cellular response. In summary we can infer that pre clinical evaluation of both MPL-A and CpG adjuvant were equally good in eliciting immune response against SARS-CoV-2 antigen and further studies are very much required to determine the actual capability of these adjuvants in developing an effctive vaccine against SARS-CoV-2.

## Acknowledgements

Authors thank DBT for the financial support under INDIGO program (GAP0872) and SS thank DST for the award of fellowships (GAP0774). Authors acknowledge Biorender software for graphical abstract illustration. IICT publication number: IICT/Pubs./2022/113. The production of soluble RBD was supported by Department of Biotechnology (DBT), Govt. of India through THSTI core grant. The following reagent was deposited by the Centers for Disease Control and Prevention and obtained through BEI Resources, NIAID, NIH: SARS Related Coronavirus 2, Isolate USA-WA1/2020, NR-52281. The following reagent was contributed by David Veesler for distribution through BEI Resources, NIAID, NIH: Vector pcDNA3.1(-) Containing the SARS-Related Coronavirus 2, Wuhan-Hu-1 Spike Glycoprotein Receptor Binding Domain (RBD), NR-52422. SARS-CoV-2 virus work was conducted inside BSL3 in the Translational Health Science and Technology Institute (THSTI), Infectious Disease Research Facility (IDRF facility).

